# The evolution of alternative splicing in *Drosophila*

**DOI:** 10.1101/054700

**Authors:** Lauren Gibilisco, Qi Zhou, Shivani Mahajan, Doris Bachtrog

## Abstract

Alternative pre-mRNA splicing (“AS”) greatly expands proteome diversity, but little is known about the evolutionary landscape of AS in *Drosophila*, and how it differs between embryonic and adult stages, or males and females. Here we study the transcriptome from several tissues and developmental stages in males and females from four species across the *Drosophila* genus. We find that 20-37% of multi-exon genes are alternatively spliced. While males generally express a larger number of genes, AS is more prevalent in females, suggesting that the sexes adopt different expression strategies for their specialized function. While the number of total genes expressed increases during early embryonic development, the proportion of expressed genes that are alternatively spliced is highest in the very early embryo, before the onset of zygotic transcription. This indicates that females deposit a diversity of isoforms into the egg, consistent with abundant AS found in ovary. Cluster analysis by gene expression levels (“GE”) show mostly stage-specific clustering in embryonic samples, and tissue-specific clustering in adult tissues. Clustering embryonic stages and adult tissues based on AS profiles results in stronger species-specific clustering, and over development, samples segregate by developmental stage within species. Most sex-biased AS found in flies is due to AS in gonads, with little sex-specific splicing in somatic tissues.

## INTRODUCTION

Alternative pre-mRNA splicing (“AS”) greatly expands the proteome diversity within species by creating different combinations of exons from the same genomic loci (Nilsen and Graveley 2010). The resulting mRNA isoforms are usually expressed in a tissue or developmental-stage specific manner, and underlie numerous essential biological processes like sex determination (Cline and and Meyer 1996; Schutt and Nothiger 2000), tissue development (Wang et al. 2008), and stress response (Staiger and Brown 2013). AS can also greatly increase the proteome diversity between species with similar repertoires of protein-coding genes (Barbosa-Morais et al. 2012; Merkin et al. 2012). Given the significance of alternative splicing in the evolution of organismal complexity (Nilsen and Graveley 2010), its dynamics across developmental stages, tissues, and species has attracted great attention (Barbosa-Morais et al. 2012; Merkin et al. 2012). Recent comparisons of transcriptomes of equivalent adult organs in several vertebrate species revealed that AS differs significantly in its complexity across the studied tetrapods, with primates showing the highest abundance of AS events (Barbosa-Morais et al. 2012; Merkin et al. 2012). Interestingly, AS shows a greater level of interspecific divergence and lineage-specific turnover across tissues than absolute gene expression levels, suggesting that the diversification of splicing significantly contributes to lineage-specific adaptation (Barbosa-Morais et al. 2012; Merkin et al. 2012).

AS is also prevalent in *Drosophila*, and has been most extensively studied in *D. melanogaster* (Chang et al. 2011; Hartmann et al. 2011; Telonis-Scott et al. 2009; Gan et al. 2010; Brooks et al. 2011; Cherbas et al. 2011; Daines et al. 2011; Venables et al. 2012; McIntyre et al. 2006). Over half of the spliced *D. melanogaster* genes encode two or more transcript isoforms, with about 50 genes capable of encoding over 1000 transcript isoforms each (Brown et al. 2014). The complexity of AS also differs across developmental stages and tissues (Graveley et al. 2011; Duff et al. 2015) and different environmental perturbations (Brown et al. 2014). However, the spatial and temporal evolution of AS between *Drosophila* species remains to be explored. Here we study the transcriptome from several tissues and developmental stages in males and females from four *Drosophila* species, spanning a major phylogenetic range of *Drosophila*. In particular, we analyze two species pairs that diverged from each other at varying evolutionary distances: *D. pseudoobscura* and its sister species *D. miranda* split about 2 million years (MY) ago (Bachtrog and Charlesworth 2002), and they diverged from *D. melanogaster* roughly 25 MY ago (Robinson 2012); *D. nasuta* and *D. albomicans* split only about 0.1 MY ago (Bachtrog 2006), and diverged from *D. melanogaster* over 60 MY ago (Tamura et al. 2004). This allows us to address novel aspects of transcriptome diversity, such as the evolutionary landscape of AS in *Drosophila*, and how it differs between embryonic and adult stages, and males and females.

## RESULTS

### Transcriptome diversity across species

We generated RNA-seq data for different tissues in *D. pseudoobscura*, *D. miranda*, *D. albomicans*, and *D. nasuta* from males and females (one female and one male larval stage and five female and five male adult samples). In addition, we used published data from sexed embryonic stages (eight female and eight male embryonic stages) in *D. pseudoobscura* and *D. miranda* (Lott et al. 2014). We analyzed a total of 144 samples from different tissues or developmental stages, summing up to a total of 6,453,796,999 reads. An overview of the data used for each species is displayed in Table 1.

**Table 1.**
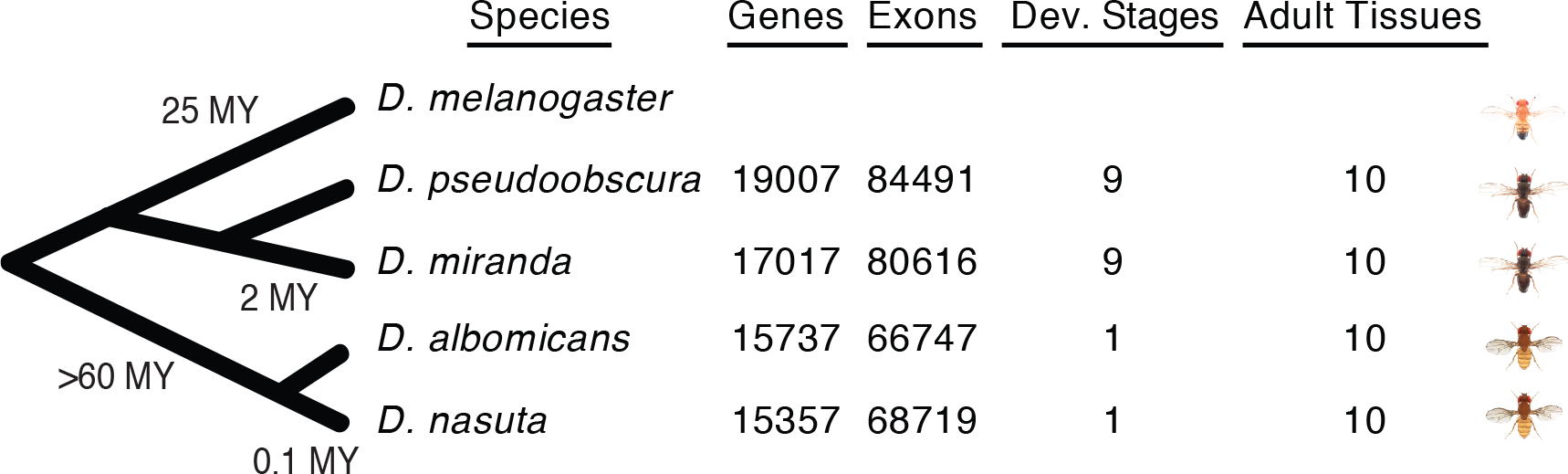
Overview of RNA-seq data used for analyses. Shown are number of genes and exons detected and numbers of stages/tissues for each of four *Drosophila* species.

The large number of stages and tissues allowed us to comprehensively annotate the transcriptomes of the four species, and a summary of the gene annotation and alternative splicing events is outlined in Figure 1. We considered skipped exons, alternative 5’ and 3’ spliced sites, mutually exclusive exons, and retained introns. We annotated between 15,357 and 19,007 genes, and between 66,747 and 84,491 exons for each species (Table 1). We find that between 20 and 37% of all multi-exon genes are alternatively spliced in at least one tissue or stage and between 5,084 and 10,172 exons (8-12% of all exons) are alternatively spliced (Table 1 and Figure 1). These values are similar to reports in *D. melanogaster* (where 31% of genes were found to be alternatively spliced; Daines et al. 2011), but considerably lower than the degree of AS in mammals, where almost all multi-exon genes are alternatively spliced (Merkin et al. 2012).

**Figure 1.**
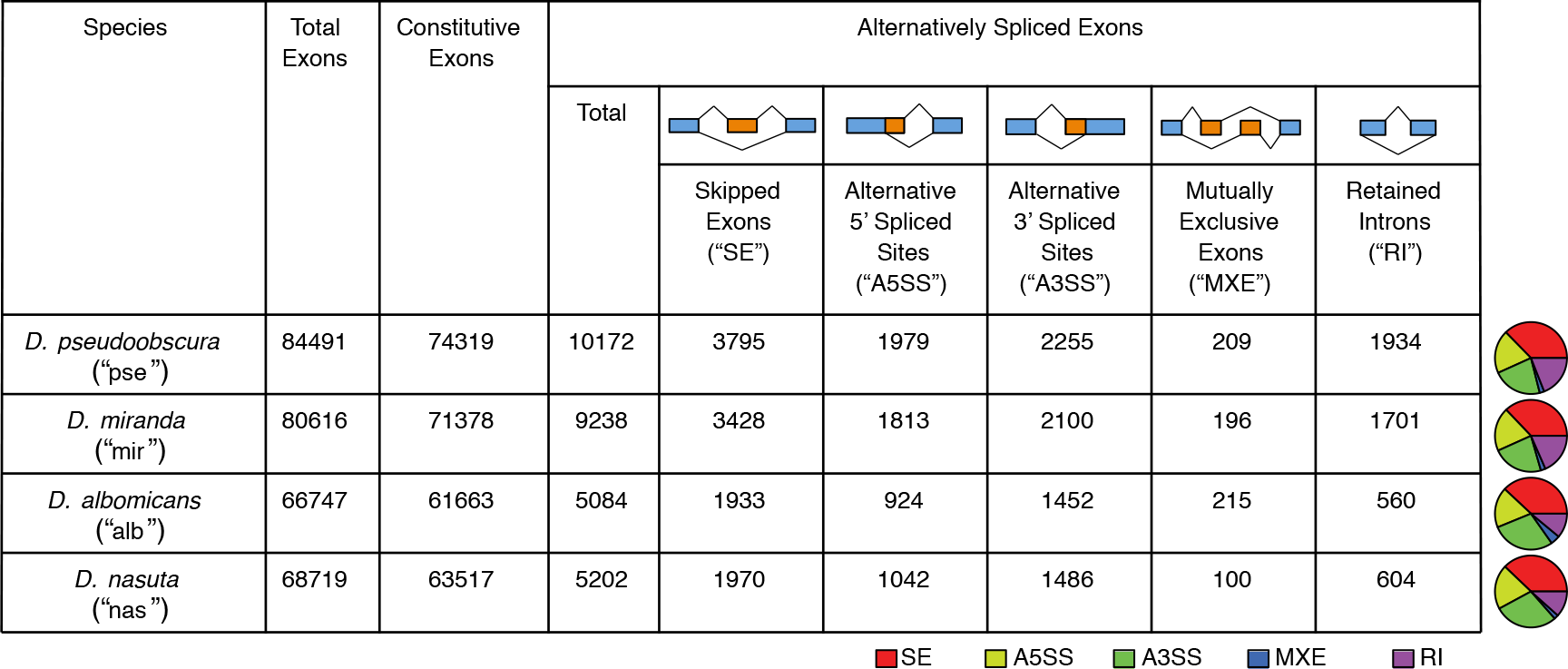
Overview of alternative splicing in four *Drosophila* species. Shown are numbers of total, constitutive, and alternatively spliced exons, as well as a breakdown of types of alternatively spliced exons, for each species. Proportions of different types of alternatively spliced exons are shown as pie charts to the right of the table.

As expected, we detected more AS features for the species for which we had more comprehensive sampling (*D. pseudoobscura* and *D. miranda*). Skipped exons are the most abundant AS event in all four species, and mutually exclusive exons are the least abundant, consistent with previous studies in *D. melanogaster* (Daines et al. 2011). The relative percentage of each type of AS is generally similar between the studied *Drosophila* species (Figure 1), despite their evolutionary distance or different numbers of tissues and developmental stages sampled, indicating a high level of conservation of AS composition.

### Temporal and spatial expression dynamics among species

We examined the transcriptome composition across corresponding samples of the four species using both the abundance of expressed genes and abundance of different alternatively spliced exons. Consistent with previous findings (Zhang et al. 2007), males generally express more genes than females across almost all tissues (Figure 2A, left panel). For example, 63-80% of all genes are expressed in male whole body while only 47-64% of all genes are expressed in female whole body. The largest discrepancy in the number of expressed genes between sexes is found in adult gonads, with testis expressing 1.6-1.7x more genes than ovary (58-76% of all genes are expressed in testis while only 35-47% of all genes are expressed in ovary). That, however, does not necessarily mean ovary or female tissues have lower transcriptome diversity. While female samples usually show fewer expressed genes, the fraction of expressed genes that are alternatively spliced is generally higher in females than in males (Figure 2A, right panel). Ovary and spermatheca have the highest percentage of expressed genes annotated as alternatively spliced (Figure 2A, right panel) despite showing among the lowest percentage of expressed genes. This indicates that male and female reproductive tissues may adopt different expression strategies for their specialized function: male tissues increase their transcriptome diversity by expressing different types of genes, while female tissues rely more on AS to increase the number of transcripts.

**Figure 2.**
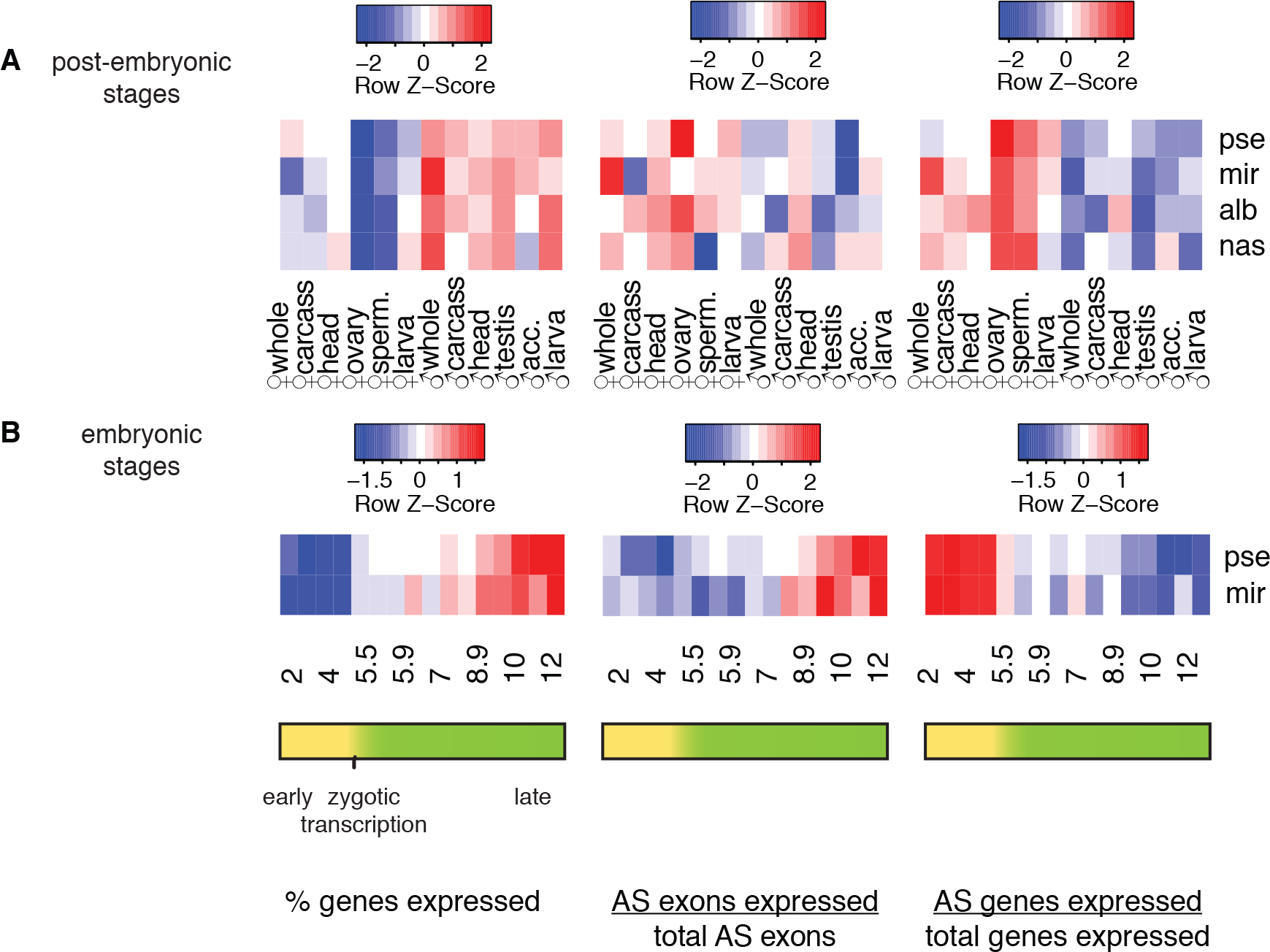
Comparisons of % of total genes expressed (left panel), % annotated alternatively spliced exons expressed (middle panel), and % of total genes expressed that are annotated as alternatively spliced (right panel) for (A) post-embryonic tissues and (B) embryonic stages. Each row of each heatmap is scaled separately by Z-score. ‘pse” = *D. pseudoobscura*; “mir” = *D. miranda*; “alb” = *D. albomicans*, “nst” = *D. nasuta* “sperm.” = spermatheca, “larva” = 3^rd^ instar larva, “acc.” = accessory gland

Between 54-71% of all genes are expressed in head, and head generally shows relatively high proportions of all annotated alternative exons expressed (8-13%; Figure 2A, middle panel). This is consistent with earlier studies of vertebrates in which brain was shown to exhibit exceptionally high transcriptional diversity (Mohr and Hartmann 2014).

Over development, both the percentage of total genes expressed (Figure 2B, left panel) and the percentage of alternatively spliced exons expressed (Figure 2B, middle panel) increase as development proceeds in both males and females. However, the proportion of expressed genes that are alternatively spliced is the highest in early embryonic development, before the onset of zygotic transcription, and drops in later stages as single-transcript gene expression increases (Figure 2B, right panel). This indicates that the mother deposits a diversity of isoforms into the egg and that early zygotic transcription increases the number of genes expressed, but most of those genes do not encode multiple isoforms. Many genes annotated in *D. melanogaster* as maternally deposited (vs. zygotic expressed or both maternal and zygotic; Lott et al. 2011), such as *bbc*, a phosphotransferase, and *Dhc64C*, a gene with ATPase activity, have multiple maternally-deposited isoforms, as alternative splicing is detected before zygotic expression begins in embryos of *D. pseudoobscura*, and *D. miranda*.

Of maternal, zygotic, and both maternal and zygotic genes defined in *D. melanogaster* (Lott et al. 2011) for which we recovered orthologs in all four species, zygotic genes show the lowest proportion of AS. However, genes annotated as being in one category in *D. melanogaster* are not necessarily in the same category in other species (e.g. a maternal gene in *D. melanogaster* may be maternal + zygotic in *D. pseudoobscura*). We therefore simply distinguished between genes that are maternally deposited (and that may or may not also show zygotic expression) as genes expressed at developmental stage 2 and zygotic genes (genes not expressed at stage 2 but expressed at later stages of embryonic development) for which we recovered orthologs in *D. pseudoobscura*, and *D. miranda*. Maternally deposited genes show a higher proportion of AS than zygotic genes in both species (*D. miranda*: 180 of 2173 (8.3%) maternally deposited genes are alternatively spliced vs. 7 of 647 (1.1%) zygotic genes alternatively spliced; *D. pseudoobscura*: 159 of 2149 (7.4%) maternally deposited genes alternatively spliced vs. 2 of 704 (0.3%) zygotic genes alternatively spliced). Thus, maternally deposited mRNA may comprise higher transcriptome diversity than previously appreciated, consistent with the abundance of alternatively spliced transcripts that we detect in ovaries. Among the zygotic genes, we confirmed sex-specific alternative splicing events of the *Sxl* gene, the master sex determining gene of *Drosophila*, during early embryogenesis in *D. pseudoobscura* and *D. miranda* (see below).

Gene ontology analysis using GOrilla (Eden et al. 2007; Eden et al. 2009) revealed enriched categories of alternatively spliced genes for the four species. Noteworthy enriched categories in genes alternatively spliced in adults include axon guidance, neuron recognition, axonal defasciculation, synaptic transmission, and learning or memory, consistent with many AS events detected in head, as well as processes related to regulation and homeostasis (**Supplementary Table 3**). Genes that are alternatively spliced during embryonic development are enriched for, among other categories, processes related to axons and neurons, synaptic processes, eye development and morphogenesis, learning, memory, behavior, and sensory organ development. Embryonic alternatively spliced genes are also enriched in processes related to reproduction, female gamete generation, and germ cell development, as well as regulation and homeostasis, development and morphogenesis, and the immune system (**Supplementary Table 4**).

### Gene expression and alternative splicing across species & development

We compared gene expression levels and alternative splicing profiles across stages (*D. pseudoobscura* and *D. miranda*) and tissues (*D. pseudoobscura*, *D. miranda*, *D. albomicans* and *D. nasuta*). As a measure of gene expression for a gene, we used TPM values (<u>T</u>ranscripts <u>P</u>er <u>M</u>illion), while alternative splicing was quantified using Ψ (<u>P</u>ercent of exons <u>S</u>pliced <u>I</u>n/“PSI”), the proportion of isoforms containing an alternatively spliced exon. When clustering adult samples on the basis of how gene expression levels correlate in pairwise comparisons, there is no strong species-specific clustering, and clustering is almost completely tissue-specific (Figure 3A). Clustering embryonic stages based on gene expression levels (Figure 3B) results in stage-specific clustering across embryogenesis, and within broad groups of stages, species-specific clustering. Gene expression during embryogenesis is conserved over particular groups of stages, and in adult stages, tissue-specific expression patterns dominate.

We looked at gene expression over embryogenesis for two subsets of genes: maternally deposited genes (which may or may not also show zygotic transcription), and zygotic genes, restricting our analysis to genes that fell into the same category in both *D. pseudoobscura* and *D. miranda*. Interestingly, when looking at expression of only zygotic genes, we see slightly more stage-specific clustering (stages 10 and 12 cluster by stage more than by species) than when we look at maternal genes that show species-specific clustering (**Supplementary Figure 1**). This suggests that expression of genes with maternally deposited transcripts is less conserved between species than genes that are expressed by the zygote during early development. Together with the observation that zygotic genes show a lower proportion of alternative splicing, this supports the idea that AS is a major source of species-specific differences and therefore is not as prevalent in conserved processes such as early embryonic development.

**Figure 3.**
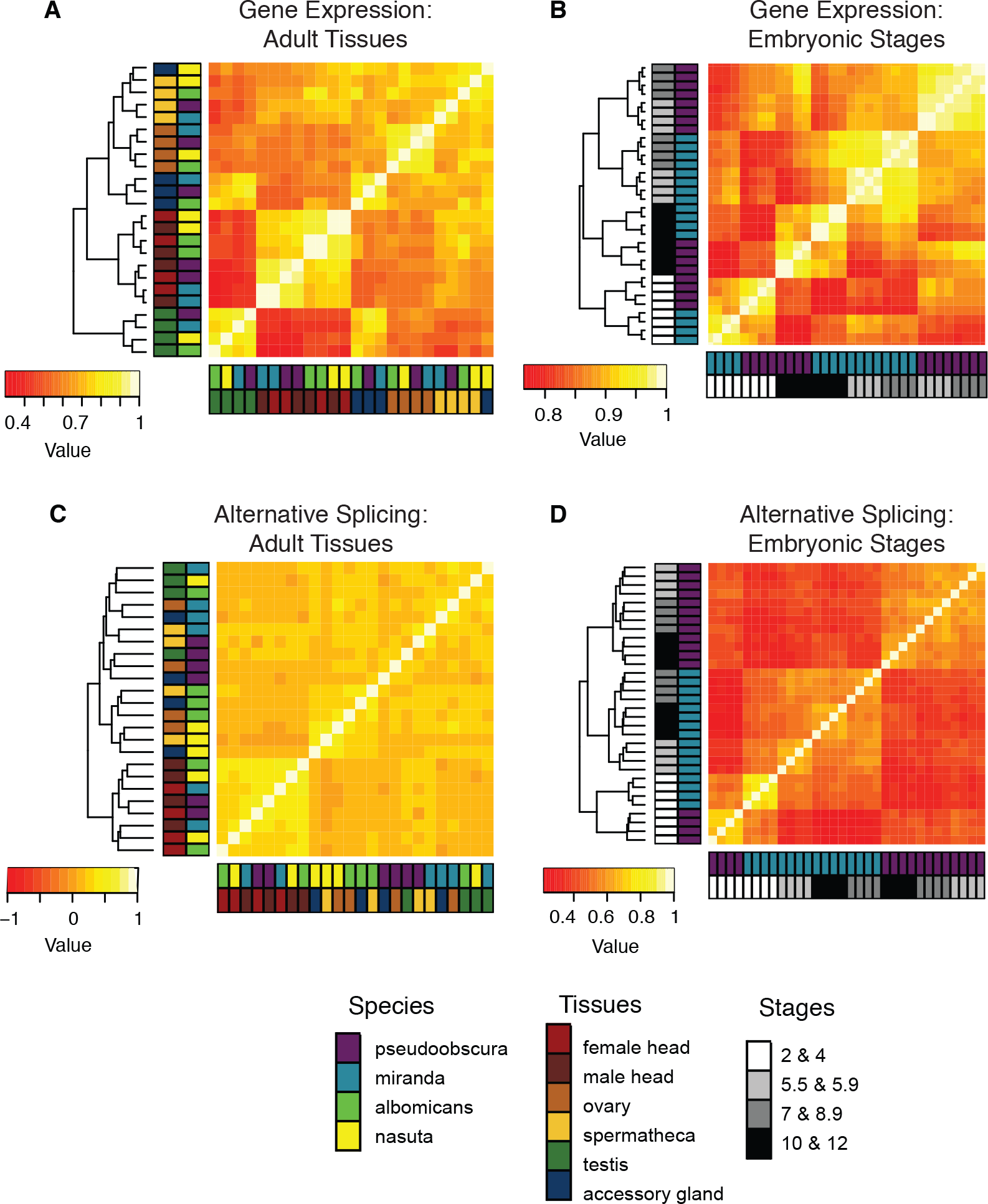
Spearman correlations based on gene expression (TPM) for genes orthologous in (A) adult tissues (n=3005) and (B) embryonic stages (n=6707) (mean of three replicates per sex/stage). Spearman correlations based on alternative splicing (Ψ) for exons orthologous and annotated as alternatively spliced in at least one sample in (C) adult tissues (n=472) and (D) embryonic stages (n=1122).

When clustering alternative splicing profiles of adult tissues using Ψ, species-based clustering tends to be stronger than tissue-based clustering (Figure 3C). Embryonic samples segregate by developmental stage (Figure 3D): prezygotic stages (2 and 4) and postzygotic stages (5.5 – 12) cluster, and within those clusters, the samples cluster by species. The more tissue-based clustering in adults observed for gene expression and the stronger species-based clustering in all samples based on splicing (for gene expression, stages 2 & 4 and stages 10 & 12 cluster by stage, while only stages 2 & 4 cluster by stage for splicing) is consistent with the strong species-specific clustering observed for splicing and “tissue-dominated clustering” observed for gene expression among vertebrates (Merkin et al. 2012). The degree of clustering by species of embryonic stages suggests that, unlike for adult samples, early development has sufficiently diverged between *Drosophila* species, both in terms of absolute gene expression, as well as based on levels of alternative splicing profiles. A comparison between gene expression and splicing has not yet been done over embryonic development in vertebrates, and it will be of great interest to see if embryogenesis in vertebrates may also be more divergent among species than adult tissues.

PCA analysis of splicing over embryonic development shows that different embryonic stages have distinct splicing profiles, which form a clock-like pattern in the PCA plot corresponding to the developmental time course (Figure 4; shown is *D. pseudoobscura*, and similar trends are seen for the other species; see **Supplementary Figures 3-5**). This differs from gene expression, which does not differentiate the different embryonic stages to the same degree as splicing. Interestingly, PCAs based on AS profiles (but not GE profiles) show that ovary is the closest of all post-embryonic tissues to embryonic stages (Figure 4). This suggests that maternally deposited alternatively spliced mRNAs play an important role in early embryonic development.

**Figure 4.**
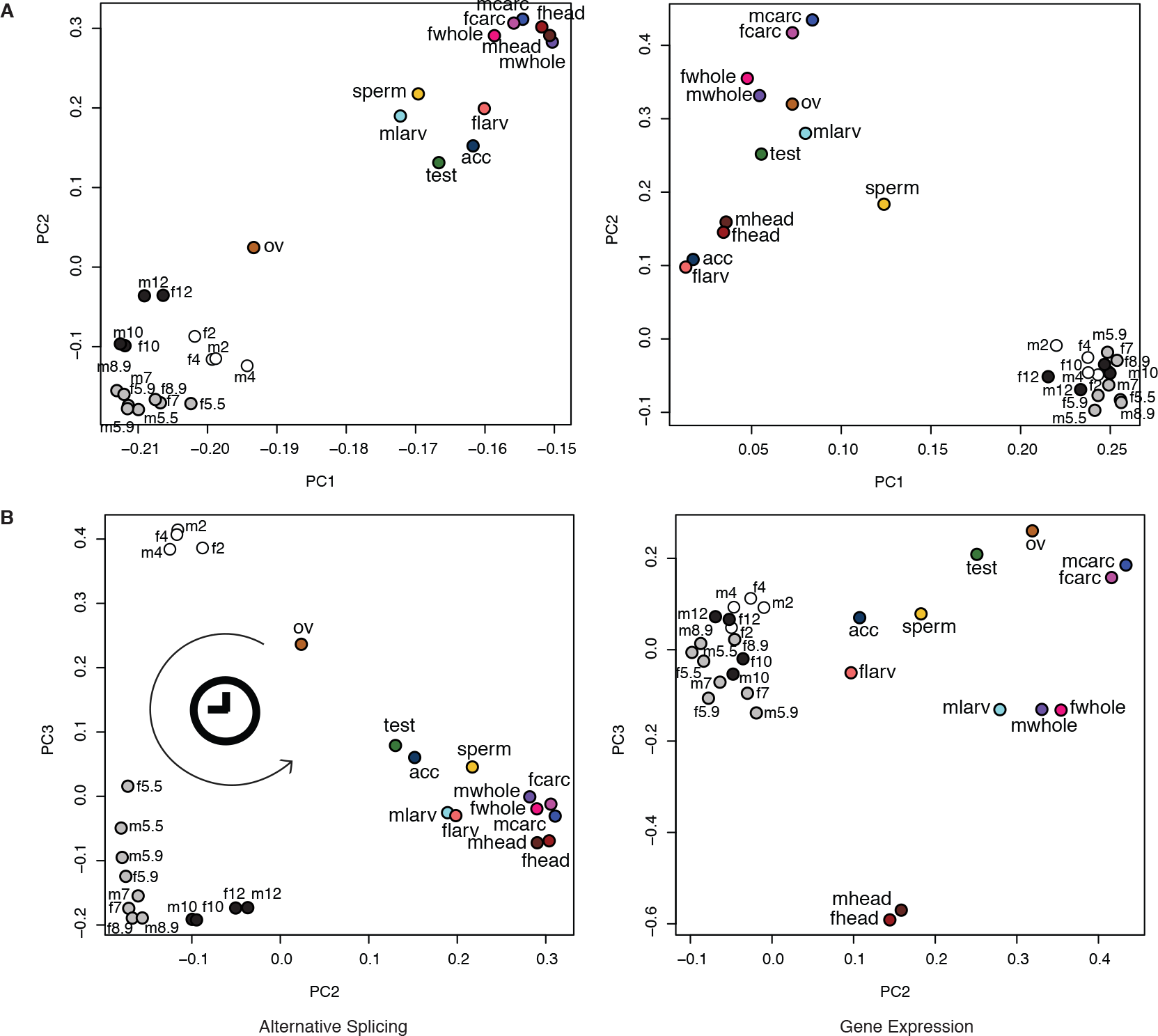
Alternative splicing (left column) and gene expression (right column) profiles for *D. pseudoobscura*. The R function *prcomp* was used to perform the PCAs. (A) PC1 (AS: 57.9% of the variance & GE: 50.1% of the variance) and PC2 (AS: 10.7% of the variance & GE: 12.4% of the variance). (B) PC2 and PC3 (AS: 4.5% of the variance & GE: 7.5% of the variance).

### Sex-biased and sex-specific alternative splicing

Alternative splicing mediates sex determination in Drosophila and our RNA-Seq data confirm sex-biased splicing of genes involved in sex determination. For example, we detect one of the exons included in males and spliced out in females for *Sxl* in the two *Drosophila* species for which we have developmental expression data (**Supplementary Figure 6**). We used ΔΨ values (ΔΨ = |Ψ_female_ − Ψ_male_|) to assess sex-biased alternative splicing in various male and female tissues and stages (Figure 5). We find sex-biased exons from genes previously observed to have sex-biased isoforms (Chang et al. 2011), such as the male-biased isoform of *thin*, a gene involved in protein ubiquitination. Proportionally, gonads show more pronounced sex-biased splicing, and the splicing pattern for gonadectomized whole flies is skewed towards weaker sex-biased splicing, for all species (Figure 5). For example, in *D. pseudoobscura* carcasses, 8.5% of AS exons are strongly sex-biased (ΔΨ>0.7) while in *D. pseudoobscura* gonads (ovary and testis), 14.8% of AS exons are strongly sex-biased (see **Supplementary Figures 8-10** for ΔΨ analyses for other species). As mentioned, heads show the highest number of AS events, but most of them show only very weak sex-bias (ΔΨ < 0.10). For example, in *D. pseudoobscura* heads, 43% of sex-biased exons have a ΔΨ < 0.10, while in gonads only 26% of sex-biased exons have a ΔΨ < 0.10. These trends are also observed in the other species analyzed (see **Supplementary Materials**). Thus, most of the sex-biased AS found in flies can be attributed to AS in gonads.

**Figure 5.**
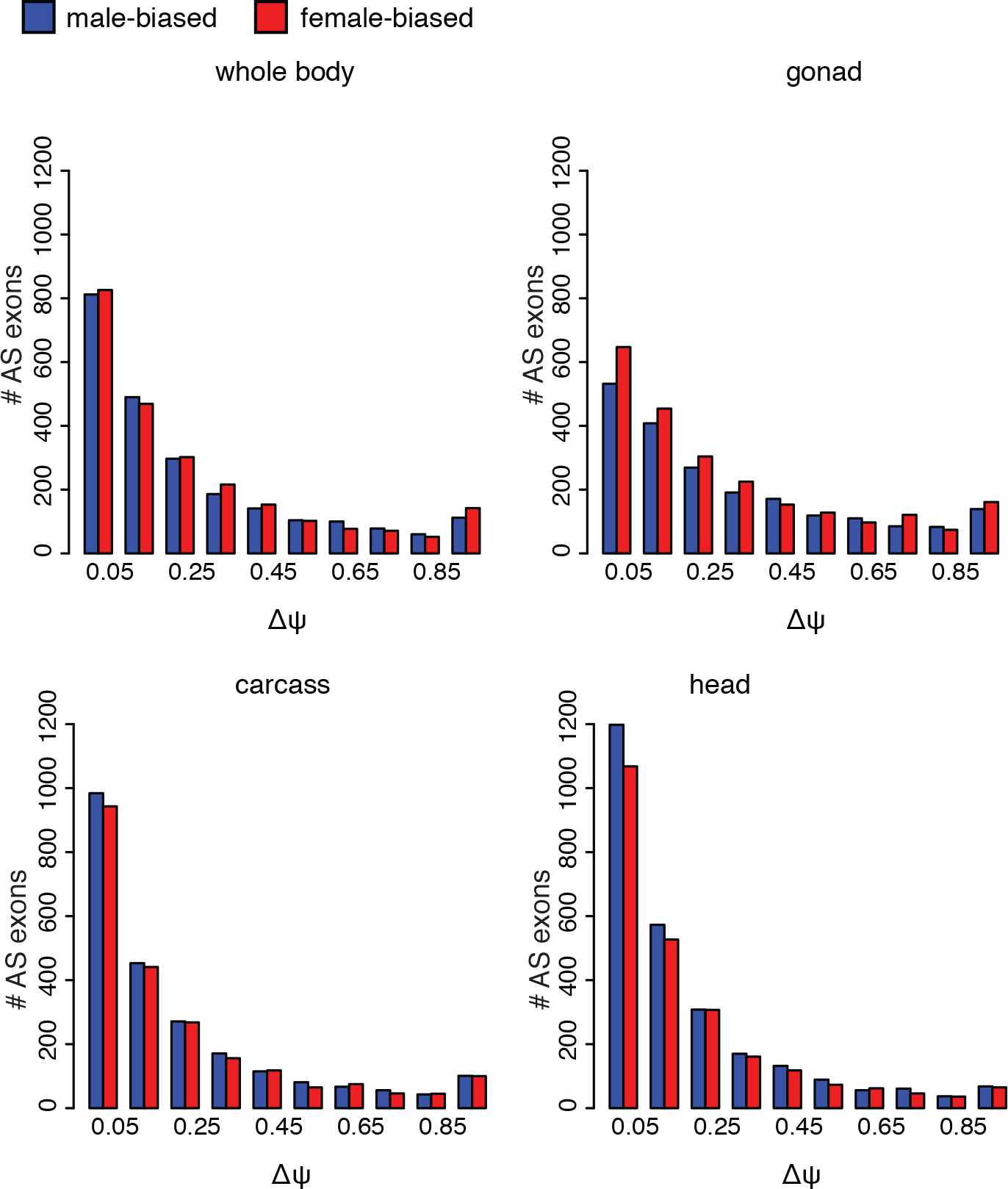
Sex-biased splicing in *D. pseudoobscura* as described by ΔΨ distributions. Comparisons are between males and females for whole body (top left), gonad (ovary and testis, top right), carcass (bottom left), and head (bottom right). The x-axis gives ΔΨ values and the y-axis shows the number of sex-biased exons. Red bars represent female-biased exons (Ψ_female_ – Ψ_male_ >0) and blue bars represent male-based exons
(Ψ_male_ – Ψ_female_ >0).

## DISCUSSION

Consistent with previous studies, we find that AS significantly contributes to increasing the transcriptome diversity in all *Drosophila* species examined, and approximately 40% of genes are alternatively spliced. In every species studied, we find that head tissue harbors the largest number of alternatively spliced exons, and the highest fraction of expressed genes that are alternatively spliced is found in ovary. This is supported by gene ontology analysis showing that alternatively spliced genes are enriched in reproduction, behavior, and neurologic processes. While males generally show a higher number of genes expressed than females, female-specific tissues (ovary and spermatheca) have the highest percentage of alternatively spliced genes. Thus, male tissues may increase their transcriptome diversity by expressing more genes, while female tissues increase the number of transcripts through AS.

Gene expression levels have been found to cluster by tissue across different mammalian species. Our analysis of Drosophila adult tissues reveals similar strong tissue-specific clustering, with only one exception (*D. nasuta* accessory gland clustering with *D. nasuta* spermatheca). This is interesting because the mammalian studies mostly examined somatic tissues (such as liver or kidney) with the exception of male testis. In contrast, most of our samples (with the exception of male and female head) contain gonad tissue (ovary/testis) or sex-specific tissues (spermatheca/accessory gland), which have been shown to be rapidly evolving (Grassa and Kulathinal 2011; Prokupek et al. 2008; Swanson and Vacquier 2002; Begun et al. 2000; Tsaur and Wu 1997). However, while sex-specific tissues may evolve more rapidly than somatic tissues, our analyses show that gene expression profiles of those tissues are conserved among species.

Similar to the findings in mammals, we see strong species-specific clustering for alternative splicing in *Drosophila*. One caveat is that the one purely somatic tissue, head, clusters by tissue. This could mean that AS in gonad tissue and sex-specific tissues may be evolving more rapidly, and AS may be more conserved in somatic tissue. Our interspecies AS analyses focused on all exons annotated as alternatively spliced in at least one sample (for adult tissue splicing analyses, n = 472 exons). However, we only recovered 49 exons alternatively spliced in at least one sample in more than one species. This implies that many exons in flies that are alternatively spliced in one species are constitutive in all the others. Species-specific splicing differences in *Drosophila* therefore may be based mostly on the binary category of whether an exon is alternatively spliced or constitutive rather than on differences in Ψ values of orthologous exons among species.

While species-specific clustering for alternative splicing is consistent with lineage-specific adaptive evolution (Barbosa-Morais et al. 2012; Merkin et al. 2012) it may also support the hypothesis that much splicing is due to erroneous splice site choice, producing non-functional isoforms targeted for degradation/NMD (Pickrell et al. 2010; Lareau et al. 2007). These presumably deleterious splicing events would therefore be unlikely to be evolutionarily conserved among species.

We find that tissue differences in gene expression patterns in adults are more dominant than sex differences; for example, male heads are closer to female heads than other male tissues. We also find evidence for considerable sex-specific splicing; however, most of the extreme differences in splicing between sexes are due to differences in splicing profiles in gonads.

In the middle of embryonic development (stages 5.5 – 8.9), stages cluster by species based both on gene expression and alternative splicing. After embryogenesis, gene expression becomes more conserved among tissues and species-specific clustering is less dominant. Alternative splicing profiles cluster samples by species rather than tissue (with the exception of head) into adulthood, suggesting that alternative splicing has diverged more than gene expression levels among *Drosophila* species. It will be of great interest to investigate gene expression profiles during embryogenesis in vertebrates to see if similar species-specific patterns exist during early development.

## METHODS

### Data

The accession numbers for the previously publicly accessible RNA-seq datasets are listed in **Supplementary Table 5**.

The remaining samples are RNA-seq datasets generated by us from *D. pseudoobscura* (male and female 3^rd^ instar larva, spermatheca), *D. miranda* (male and female head, spermatheca), *D. albomicans* (male and female whole body, virgin male and female gonadectomized carcass, male and female head, ovary, spermatheca, testis, accessory gland, male and female 3^rd^ instar larva) and *D. nasuta* (virgin male and female whole body, male and female carcass, male and female head, ovary, spermatheca, testis, accessory gland, male and female 3^rd^ instar larva). We extracted total RNA from whole body or dissected tissues (Qiagen) and prepared RNA-seq libraries following the standard Illumina protocol. Briefly, we used Dynal oligo(dT) beads (Invitrogen) to isolate poly(A) mRNA from the total RNA samples. We then fragmented the mRNA by using the RNA fragmentation kit from Ambion, followed by first- and second-strand cDNA synthesis using random hexamer primers (Invitrogen). We complemented the cDNA synthesis by an end repair reaction using T4 DNA polymerase and Klenow DNA polymerase for 30 min at 20°C. We then added a single A base to the cDNA molecules by using 3'-to-5' exo-nuclease, and ligated the Illumina adapter. The fragments were subjected to size selection on a 2% gel and purification (Qiagen). We finally amplified the cDNA fragments by PCR reaction and examined the libraries by Bioanalyzer (Aglient). Paired-end cDNA sequencing was performed on the Illumina HiSeq 2000 at the Vincent J. Coates Genomics Sequencing Laboratory at the University of California, Berkeley. This work used the Vincent J. Coates Genomics Sequencing Laboratory at UC Berkeley, supported by NIH S10 Instrumentation Grants S10RR029668 and S10RR027303.

More information on how to access this data can be found in DATA ACCESS.

### For within-species analyses

We used seqtk (https://aur.archlinux.org/packages/seqtk-git) to randomly choose paired-end RNA-seq reads so that within each species, each sample used had the same number of reads (Supplementary Table 1b; for reads used in inter-species analyses, see Supplementary Table 1a). The pooled data from all four species had a total of 957,749,296 reads. For samples with read lengths > 50 bp, we used read quality data from FastQC (http://www.bioinformatics.bbsrc.ac.uk/projects/fastqc) to determine cutoffs and trimmed the reads to 50 bp.

We used bowtie2-build and TopHat v2.0.13 (Trapnell et al. 2009) to map the reads to each genome (*D. pseudoobscura* Release 3.1 downloaded from href="http://www.flybase.org, *D. miranda* assembly (Zhou et al. 2013), *D. albomicans* assembly (Zhou et al. 2012b), *D. nasuta* assembly, unpublished) using the parameters --b2-sensitive, --coverage-search, and --microexon-search and then ran Cufflinks v2.1.1 (Trapnell et al. 2010) without a reference annotation on each sample. For each species, we used cuffmerge to combine all of the cufflinks-generated annotations to create a master annotation. We used cuffdiff (Trapnell et al. 2013) with this master annotation to obtain expression data for each sample, using a minimum cutoff of FPKM = 1.

We used picard v1.106 (http://picard.sourceforge.net) to obtain insert size mean and standard deviation for each sample. We used MATS (Park et al. 2013; Shen et al. 2012) to get Ψ (<u>P</u>ercent <u>S</u>pliced <u>I</u>n) values for each alternatively-spliced exon by running pairwise comparisons among all samples within each species and taking an average w for each exon calculated using reads on target and junction counts, excluding samples for which the exon was not classified as alternatively spliced. To validate our pipeline, we also tested a few tissues using an alternative pipeline. We used AltEventFinder (Zhou et al. 2012a) to annotate alternative splicing events and MISO (Wang et al. 2008; Katz et al. 2010) to compute Ψ values. Supplementary Table 2 shows correlations of the results of both pipelines for post-embryonic tissues in *D. miranda* computed in R using the *corr* function.

Gene ontology analysis was performed using GOrilla (Eden et al. 2007; Eden et al. 2009) to reveal enriched categories of processes (p<10^−3^) by comparing two unranked lists of genes: target genes (alternatively spliced genes) and background genes (genes determined orthologous in all four species for adult tissues and genes determined orthologous between *D. pseudoobscura* and *D. miranda* for embryonic stages).

*heatmap.2* was used in R to generate heatmaps, which uses *hclust* to cluster samples. The R function *prcomp* was used to perform the PCAs.

### For interspecific comparison

Pairwise whole genome alignments for each pair of species were done using the software Mercator (https://www.biostat.wisc.edu/~cdewey/mercator/) (Dewey and Pachter 2006) and MAVID (Bray and Pachter 2004). First we used Mercator to build an orthology map for each pair of species. Then MAVID was used to perform global whole genome alignments. Finally, for each exon/gene in each species, the coordinates of the corresponding ortholog in the other species was determined using the sliceAlignment program (Mercator distribution).

Using pairwise alignments of *D. pseudoobscura*, *D. miranda*, *D. albomicans*, and *D. nasuta* to *D. melanogaster*, we kept genes/exons aligned with >0.5 overlap between all pairs. We kept all genes/exons with 1:1 orthology. If there are multiple genes/exons from one species that aligned to the same *D. melanogaster* gene/exon, we kept the pair with the highest overlap score. If the highest overlap score was shared between the *D. melanogaster* gene/exon and more than one gene/exon from the other species, we did not use the gene(s)/exon(s)in our analysis. If there were multiple *D. melanogaster* genes/exons that aligned to the same gene/exon from the other species, we kept the pair with the highest overlap score. If the highest overlap score was shared between one gene/exon from the other species and more than one *D. melanogaster* gene/exon, we did not use the gene(s)/exon(s) in our analysis. For comparisons over embryonic development, this left us 6707 genes with 1:1 orthologous relationships between *D. pseudoobscura* and *D. miranda* and 1122 exons with 1:1 orthologous relationships between *D. pseudoobscura* and *D. miranda* that were also annotated as alternatively spliced in at least one sample. For comparisons of post-embryonic tissues, we recovered 3005 genes with 1:1 orthologous relationships among *D. pseudoobscura*, *D. miranda*, *D. albomicans*, and *D. nasuta* and 472 exons with 1:1 orthologous relationships among the four species that were also annotated as alternatively spliced in at least one sample.

We used orthology information to create annotations for each species containing only genes orthologous among all species (*D. pseudoobscura*, *D. miranda*, *D. albomicans*, and *D. nasuta* for comparisons of post-embryonic samples; *D. pseudoobscura* and *D. miranda* for comparisons of embryonic samples) and ran kallisto (Bray et al. 2015) using these annotations. We used TPM (<u>T</u>ranscripts <u>P</u>er <u>M</u>illion) values from each sample for each gene for our interspecies gene expression analysis. We used sleuth (Bray et al. 2015) to normalize TPM values between samples and to compute Jensen-Shannon divergence between samples.

We used bowtie2-build and TopHat v2.0.13 (Trapnell et al. 2009) to map the reads to each genome: *D. pseudoobscura* Release 3.1 (downloaded from http://www.flybase.org); *D. miranda* assembly (Zhou et al. 2013); *D. albomicans* assembly (Zhou et al. 2012b); *D. nasuta* assembly (unpublished) using the parameters --b2-sensitive, --coverage-search, and --microexon-search and then ran Cufflinks v2.1.1 (Trapnell et al. 2010) using the -G parameter and the reference annotations used for orthology analysis on each sample.

For interspecies splicing analyses, we ran MATS (Park et al. 2013; Shen et al. 2012) with the bam files and the annotations used for orthology analyses. MATS compares splicing in samples pairwise, and we compared all samples within species that shared the same number of reads. Spermatheca reads from *D. nasuta* and *D.albomicans* were trimmed to 76 base pairs (the size of all other reads in these species). In each sample, we took an average Ψ for each exon calculated using reads on target and junction counts, excluding samples for which the exon was not classified as alternatively spliced. For exons alternatively spliced in some samples but not others, we looked at the expression calculated for that exon in cufflinks. If the exon had an FPKM value < 1 and the upstream or downstream exons had an FPKM value >1, the exon was assigned a Ψ value of 0. If the exon had an FPKM value >1, the exon was assigned a Ψ value of 1. If the exon had an FPKM value <1 and the upstream and downstream exons had an FPKM value <1, the exon was not assigned a Ψ value.

## Data Availability

All sequencing reads generated in this study are deposited at the National Center for Biotechnology Information Short Reads Archive (www.ncbi.nlm.nih.gov/sra) under the Bioproject accession no. [xxx].

